# A novel C57BL/6 mouse model for the study of severe *Citrobacter rodentium* infection

**DOI:** 10.64898/2026.03.12.711355

**Authors:** Kathleen G. McClanahan, Luisella Spiga, M. Blanca Piazuelo, Jennifer A. Gaddy, Wenhan Zhu, Danyvid Olivares-Villagómez

**Affiliations:** Department of Pathology, Microbiology, and Immunology, Vanderbilt University Medical Center, Nashville, TN, USA; Vanderbilt Institute for Infection, Immunology, and Inflammation, Vanderbilt University Medical Center, Nashville, TN, USA; Division of Gastroenterology, Hepatology, and Nutrition, Department of Medicine, Vanderbilt University Medical Center, Nashville, TN, USA; Center for Mucosal Inflammation and Cancer, Vanderbilt University Medical Center, Nashville, TN, USA; Department of Medicine, Division of Infectious Diseases, Vanderbilt University Medical Center, Nashville, TN, USA; Tennessee Valley Health Systems, Department of Veterans Affairs, Nashville, TN, USA; Medicine Health and Society, Vanderbilt University, Nashville, TN, USA

## Abstract

The study of human enteropathogenic and enterohemorrhagic *Escherichia coli* (EPEC and EHEC) has been limited by the inability of these pathogens to effectively colonize murine models without prior antibiotic treatment. Because it mimics key features of human EPEC and EHEC infection, *Citrobacter rodentium*, a natural mouse pathogen that colonizes the lower intestine, has become the primary model for investigating these organisms. C57BL/6 mice are most commonly used for *C. rodentium* research, however, unless they carry specific genetic mutations, they typically develop only mild disease and clear the infection within weeks. As a result, models of severe disease in genetically unmodified hosts are lacking. Here, we describe the development of a non-genetically modified C57BL/6 mouse line with an undisturbed intestinal microbiota, highly susceptible to severe *C. rodentium* infection. Early infection in these mice was marked by significantly elevated cecal bacterial burdens and tissue pathology. Immune profiling revealed broad reductions in multiple lymphoid subsets, indicating impaired early mucosal activation. Although overall cytokine expression patterns were similar between groups, ceca of susceptible mice exhibited elevated baseline and early post-infection IL-18, as well as increased G-CSF at day 1. Microbiota analyses showed broadly comparable communities with wildtype controls, with some altered groups, such as Lachnospiraceae, Prevotellaceae, Desulfovibrionaceae, and Erysipelotrichaceae. Together, these findings characterize a robust C57BL/6 model that reproducibly develops severe *C. rodentium*-induced disease. This phenotype is driven by microbiota-associated alterations and impaired early cecal immunity, providing a valuable system for studying host–microbiota interactions in enteric infections.

## Introduction

Attaching and effacing bacteria, such as enteropathogenic and enterohemorrhagic *Escherichia coli* (EPEC and EHEC, respectively), disrupt intestinal epithelial function, leading to impaired ion and water transport, and severe bloody diarrhea in humans.^1,2^ Progress in understanding host-pathogen interactions in these infections is hindered by the fact that these bacteria do not easily colonize most small animal models. *Citrobacter rodentium*, a natural murine pathogen that colonizes the distal intestine and produces attaching and effacing lesions, has become the main model for human EPEC and EHEC. This mouse bacterium has provided key insights into the mechanisms underlying the pathogenicity used by attaching and effacing bacteria.^3-9^

Disease severity following *C. rodentium* infection varies substantially by mouse strain. For example, infected C3H/HeJ quickly lose weight, develop severe disease leading to dehydration and in most animals, death. Other strains, like C57BL/6 (one of the most widely used strains due to the availability of powerful genetic tools), develop low grade colonic inflammation, limited weight loss, and eventually clear out the infection.^10^ This relative resistance has advantages for investigating pathogen’s virulence and host protective immune response without excessive damage.

However, mild disease outcomes also limits investigation into the mechanisms that drive severe disease, which constitutes a key aspect for developing preventive and therapeutical strategies against attaching and effacing pathogens. Genetically modified mouse models provide an important tool to investigate how a resistant host can become susceptible to severe infection. For example, we have shown that granzyme B-deficient mice on the C57BL/6 background exhibit marked weight loss, increased colon pathology, hunched posture, diarrhea, rectal bleeding, and death following *C. rodentium* infection, in part because of the dysregulated IL-17 expression by CD4^+^ T cells.^11^

In parallel, growing evidence indicates that the intestinal microbiota and/or its metabolites are critical determinants of susceptibility to bacterial infections and intestinal inflammation.^12-14^ However, microbiota studies usually rely on the utilization of gnotobiotic or germ-free mice, which are costly, technically demanding, and associated with important developmental and immunologic differences from conventionally housed animals. Thus, mouse models in which the microbiota is minimally altered and present a traceable phenotype, are necessary and critical tools to study microbiota-pathogen-host interactions. In this publication, we present a novel line susceptible to *C. rodentium* infection derived from wild-type C57BL/6 mice, providing a useful platform for the study of severe intestinal infection, the microbiota and inflammation.

## Results

### Development of susceptible C57BL/6 mice

We reported that C57BL/6 granzyme B-deficient mice (*Gzmb*^-/-^) developed severe *Citrobacter rodentium* infection.^15^ However, because granzyme B is known to play an important role in cytotoxicity against bacteria and attenuation of virulence factors,^16,17^ it is possible that part of the phenotype observed was due to the absence of this enzyme. In addition, *Gzmb*^*-/-*^ susceptible (S) mice became resistant to severe disease when cohoused with resistant C57BL/6 wild type mice, or when fecal matter from resistant mice was transplanted into susceptible recipients,^15^ indicating an important role for the intestinal microbiota in this phenotype. Indeed, C57BL/6 wild type mice became susceptible to severe *C. rodentium* infection when their microbiota was depleted with antibiotics and transplanted with fecal matter from susceptible mice.^15,18^ Although fecal matter transplant restores most of the microbiota lost after antibiotic treatment, residual communities may persist, expand, and dampen expected phenotypes. To circumvent this potential issue and to generate a stable, wildtype C57BL/6 mouse line susceptible to *C. rodentium* infection, we crossed a germ-free male with non-germ free, susceptible *Gzmb*^-/-^ dams. Their F1 *Gzmb*^+/-^ offspring was interbred to generate *Gzmb*^+/+^ mice. Male and female mice derived from the above interbreeding were infected with *C. rodentium* and monitored for weight change and disease signs. Both male and female mice lost weight during the course of the infection (**Figure 1A)** and developed severe signs of disease (**Figure 1B**), whereas C57BL/6 wild type (WT) mice gained weight and remained disease free during the entire course of the experiment. Susceptibility to infection has been maintained across multiple generations, indicating that the phenotype is stable. Therefore, these mice represent a reliable, non-genetically altered, C57BL/6 line susceptible to severe *C. rodentium* infection and hereafter referred to as (S)WT mice.

**Figure 1.**
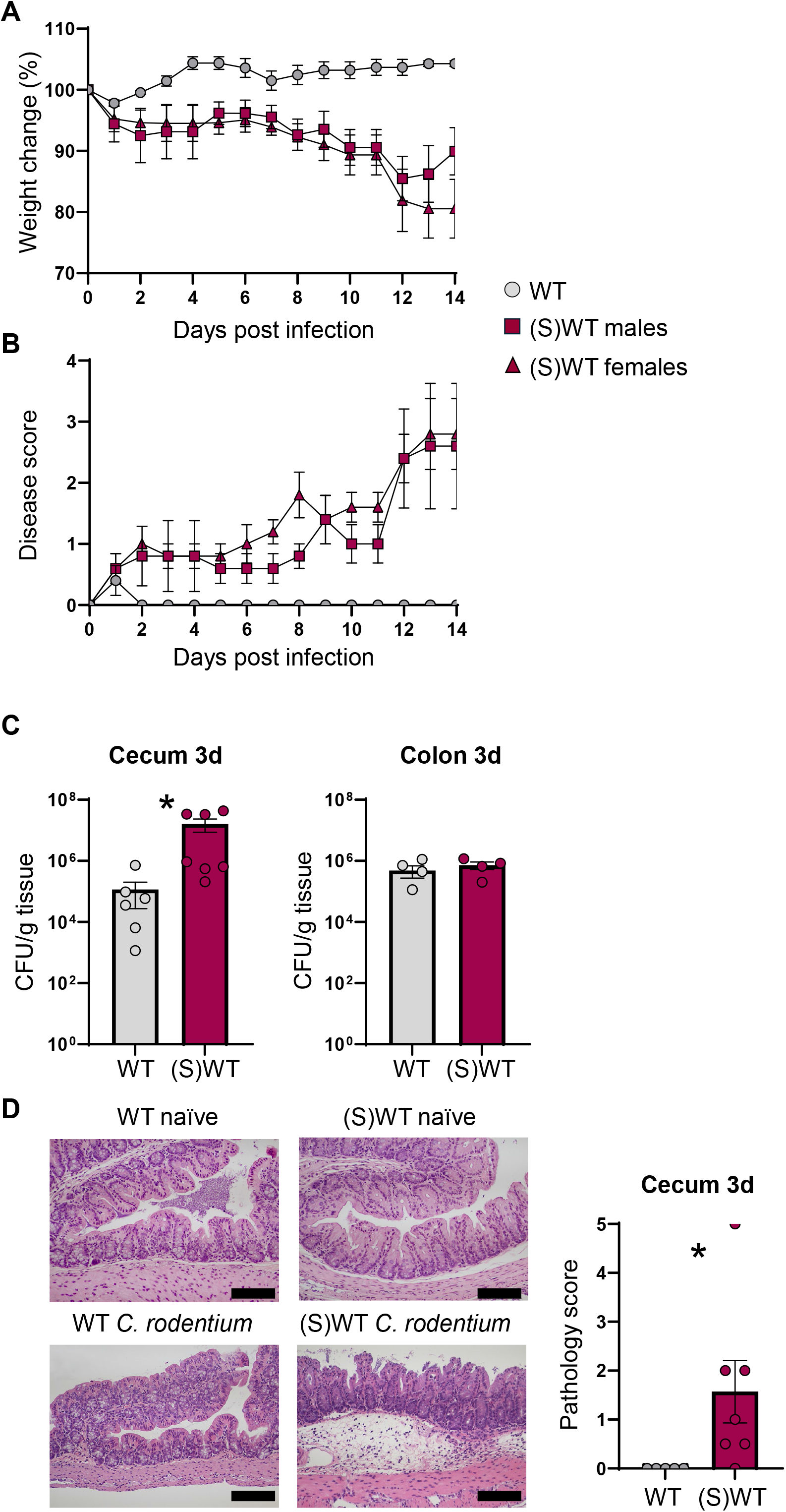
Development of C57BL/6 mice susceptible to *C. rodentium* infection. A male, germ-free mouse was bred with susceptible *Gzmb*^-/-^ females. Offspring was bred among themselves to obtain *Gzmb*^+/+^ mice, referred to as susceptible wild type mice, (S)WT. Mice were infected with 1-5×10^8^ *C. rodentium* CFU, monitored daily for weight (A) and signs of disease (B), such as diarrhea, scruffiness, rectal irritation/bleeding. WT, n = 5; (S)WT males, n = 5, (S)WT females, n = 5. (C) Cecum and colon colonization at day 3 after infection. WT cecum, n = 6; WT colon, n = 4; (S)WT cecum, n= 7; (S)WT colon, n = (D) Cecum pathology micrographs (left; bar indicates 100μm) and score (right) at day 3 after infection. WT, n = 5; (S)WT, n = 6. \**P* < 0.05, Student’s t-test. All panels are representative experiments of at least 2 replicas.

### Disease course in susceptible C57BL/6 mice

During the first three days after infection, *C. rodentium* establishes primarily in the cecum, in particular within the specialized patch of lymphoid tissue known as the cecal patch.^19,20^ As shown in **Figure 1B**, (S)WT mice developed signs of disease starting at day one post infection, including wasting, irritated rectum and loose stools. Cecum colonization at day 3 in (S)WT mice reached an average of 10^7^ colony forming units (CFU)/gram of tissue in comparison to 10^5^ CFU/g of tissue in WT mice (**Figure 1C**). In contrast, colon colonization was mostly indistinguishable at the same time point between the two groups. Increased cecum colonization correlated with higher pathological damage (**Figure 1D**).

### Cellular immune response in susceptible C57BL/6 mice

The increased *C. rodentium* cecum colonization and its associated pathology in (S)WT mice during the first days after infection, indicate a possible defect in the cecal immune response against this bacterium. To investigate this possibility, we analyzed the cellular composition in the cecal intraepithelial lymphocyte (IEL) and lamina propria (LP) compartments (as an example, **Figure 2A** shows partial flow cytometric strategy for the IEL compartment). One day after infection, the total number of CD45^+^ cells (hematopoietic origin) was higher in WT mice in comparison to (S)WT mice (**Figure 2B**), which was not observed in naïve WT and (S)WT mice (**Figure 2C**). Thus, the difference in CD45^+^ cell numbers was a result of infection and not a basal disruption, and was maintained for at least 3 days post infection (**Figure 2D**). Because CD45^+^ cells represent the majority of lymphoid cells in the IEL compartment, these results indicate a potential disruption in individual IEL populations. Indeed, when a detailed analysis of different IEL subpopulations was performed, we observed that TCRγ^+^, TCRβ^+^, TCR^neg^, iCD8α, TCRβ^+^CD8αα^+^, and TCRβ^+^CD8αβ^+^ cell numbers were significantly decreased in (S)WT mice in comparison to WT mice (**Figure 2B**).

**Figure 2.**
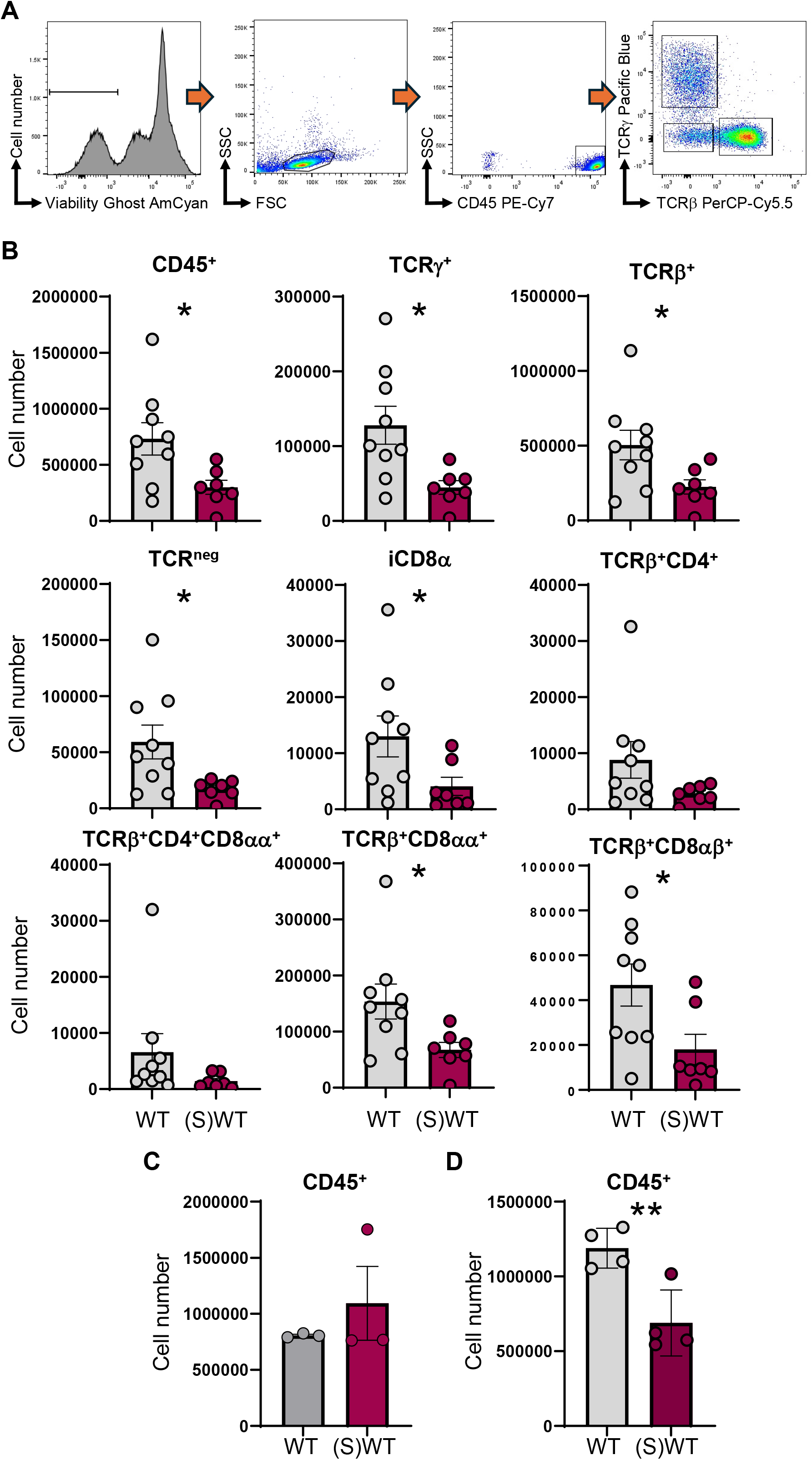
C57BL/6 mice susceptible to *C. rodentium* infection present disruption in the cecal IEL compartment. WT and (S)WT mice were infected with 1-5×10^8^ *C. rodentium* CFU. At the indicated time points, cecum IEL were isolated, counted and stained with different fluorochrome labeled antibodies. (A) Gating strategy for the IEL compartment. For clarity purposes, panel does not include gating beyond TCR usage. Orange arrows indicate gating flow. (B) Cell numbers of the indicated IEL populations at 1 day post infection. WT, n = 9; (S)WT, n = 7. (C) Total CD45^+^ cell numbers in cecal IEL derived from naïve WT (n = 3) and (S)WT mice (n = 3). (D) Total CD45^+^ cell numbers in cecal IEL at 3 days after infection. WT, n = 4; (S)WT, n = 4. \**P* < 0.05; \*\**P* < 0.01, Student’s t-test. All panels are representative experiments of at least 2 replicas.

Analysis of the cecal LP showed a more limited phenotype. With the exception of TCRβ^+^CD8αβ^+^ cells, total cell numbers of different LP populations were mostly undistinguishable between (S)WT and WT mice at 1 day post *C. rodentium* infection (**Figure 3**). Overall, these results indicate that WT mice mount an early mucosal cellular immune response to *C. rodentium*, particularly in the IEL compartment, that may restrict cecal colonization. However, (S)WT mice present a dysfunctional cellular cecal immune response that may result in higher *C. rodentium* burden and associated pathology.

**Figure 3.**
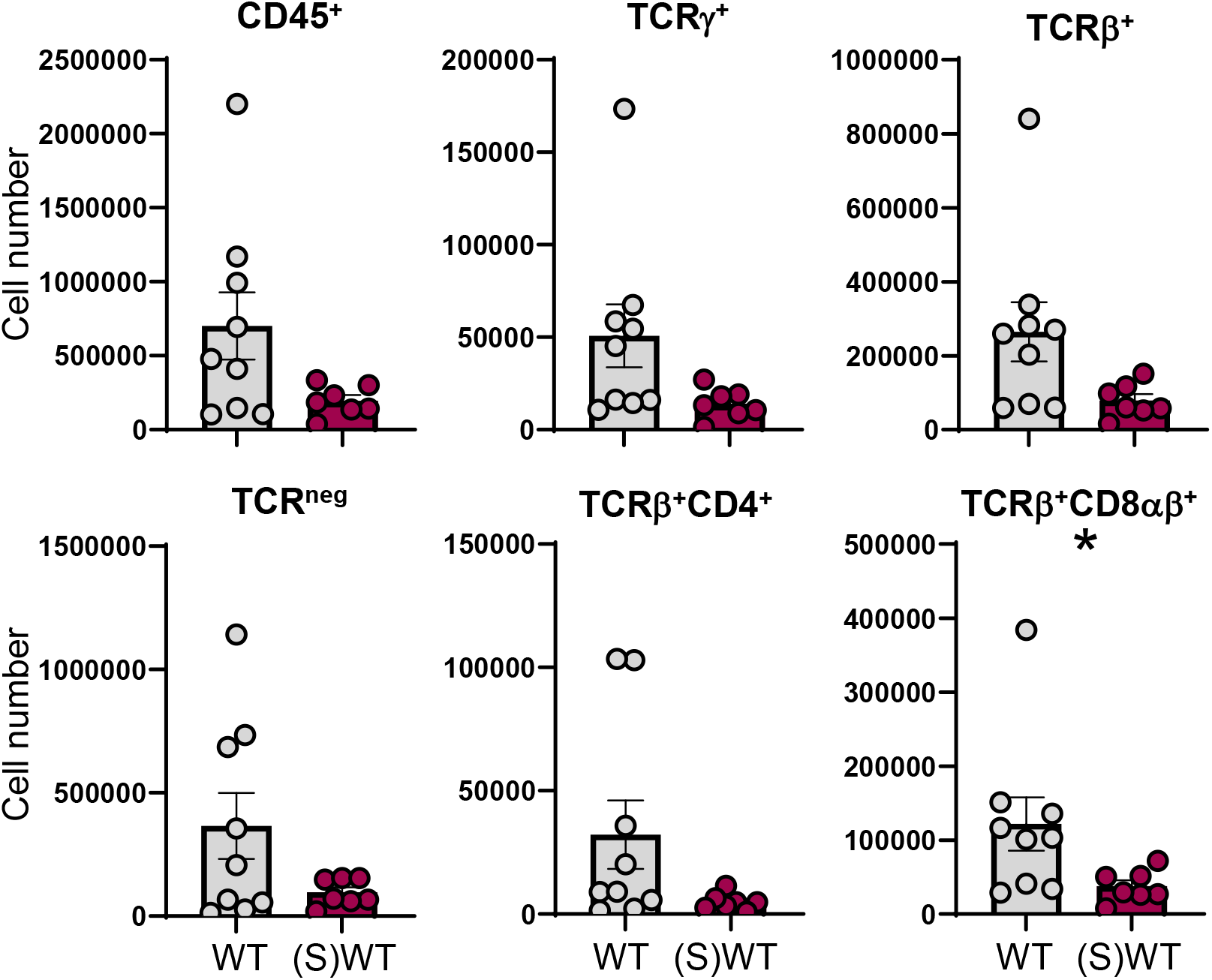
C57BL/6 mice susceptible to *C. rodentium* infection do not show an altered cecal LP immune response. WT and (S)WT mice were infected with 1-5×10^8^ *C. rodentium* CFU. At one day after infection, cecum LP cells were isolated, counted and stained with different fluorochrome labeled antibodies as in **Figure 1.** Graphs show cell numbers of the indicated populations. WT, n = 9; (S)WT, n = 7. \**P* < 0.05, Student’s t-test. All panels are representative experiments of at least 2 replicas.

### Cytokine profile in susceptible C57BL/6 mice

As reported in the previous section, we observed alterations in the IEL compartment in *C. rodentium*-infected (S)WT mice in comparison to WT counterparts. This phenotype suggests that an overall cytokine profile may also be altered in (S)WT mice. To investigate this possibility, we obtained cecal and colon from naïve, and 1- and 3-day infected mice and determined their global inflammatory cytokine profile (**Figure 4**). In naïve animals, the cytokine expression from cecum and colon appeared similar between WT and (S)WT mice, with the notable exception of higher IL-18 expression in the cecum of (S)WT mice.

**Figure 4.**
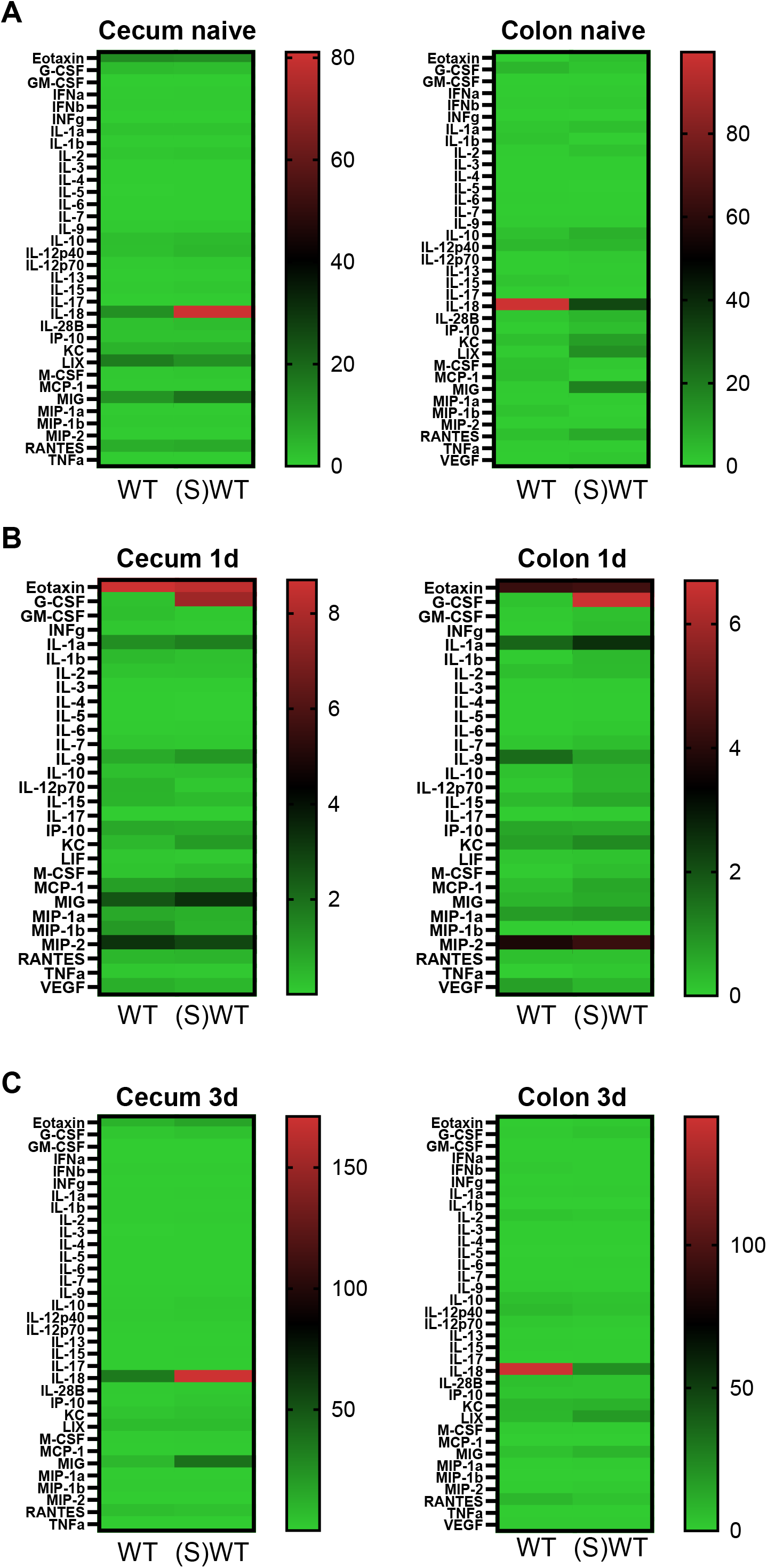
Global cytokine profile in C57BL/6 mice susceptible to *C. rodentium* infection. Luminex cytokine profile from the cecum and colon at the indicated time points after *C. rodentium* infection (1-5×10^8^ *C. rodentium* CFU per mouse). (A) Naïve mice; (B) 1 day and (C) 3 days post infection. Heatmaps show the average expression level per group. WT naïve, n = 3; (S)WT naïve, n = 4; WT 1 day, n = 4; (S)WT 1 day, n = 4; WT 3 days, n = 5; (S)WT 3 days, n = 5. Panels represent an individual experiment.

Interestingly, the opposite pattern was observed in the colon, where IL-18 expression was higher in WT mice in comparison to (S)WT animals (**Figure 4A**). One day after infection, the overall cytokine profile in the cecum and colon was similar between the two groups, except for the increased granulocyte colony stimulating factor (G-CSF) expression in (S)WT mice (**Figure 4B**). Of note, IL-18 expression at this time point was not consistently reliable. By day 3 of infection, the major difference between the two groups was increased cecal IL-18 expression in the cecum, whereas an inverted profile was observed in the colon (**Figure 4C**).

Detailed analysis of representative cytokines in the cecum showed that IL-18 was significantly increased in (S)WT mice at day 3 in comparison to WT controls (**Figure 5**).

**Figure 5.**
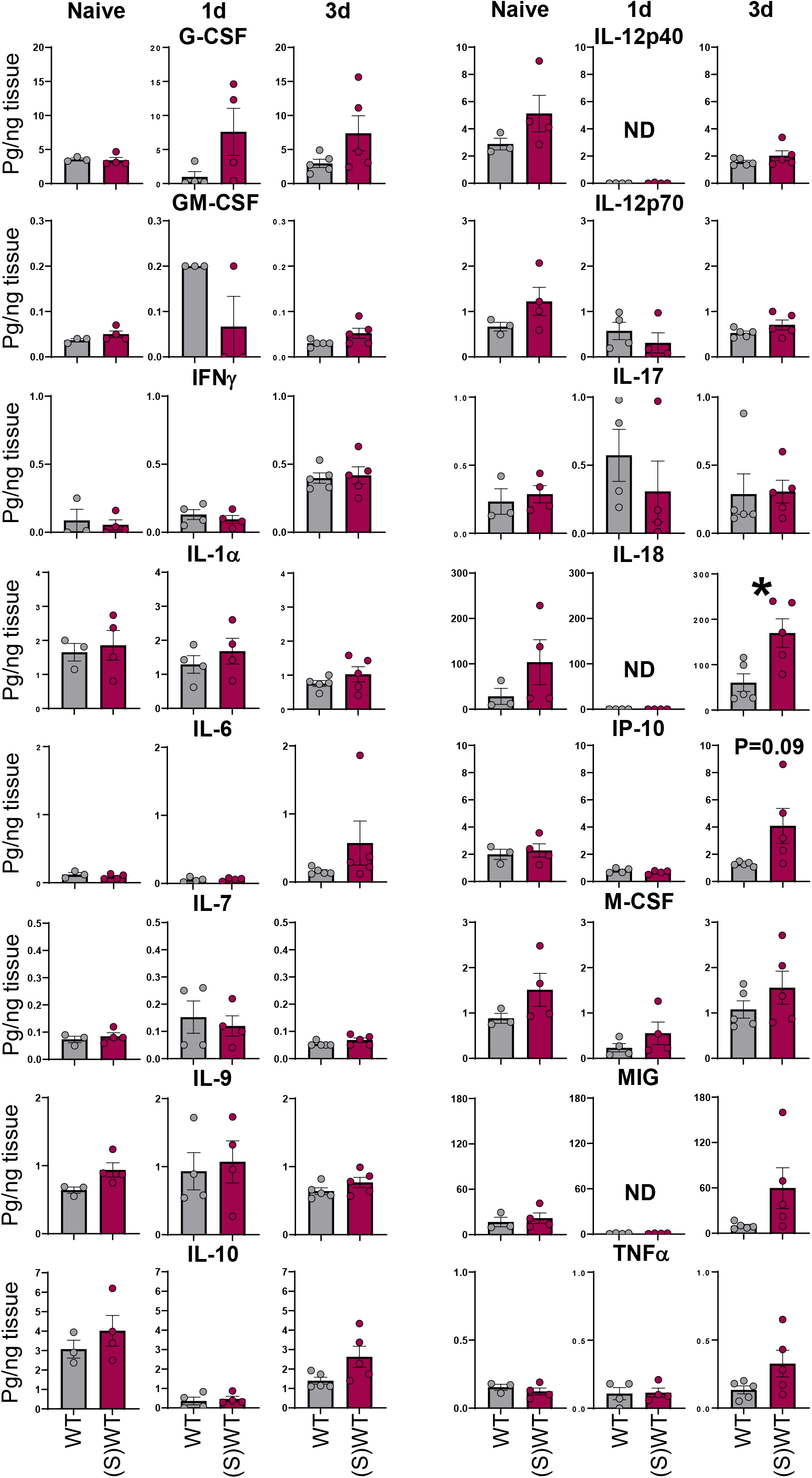
Detailed cytokine profile in the cecum of C57BL/6 mice susceptible to *C. rodentium* infection. Data derived from the experiment in **Figure 4.** Individual cytokine expression in the cecum of WT and (S)WT mice at the indicated time points after *C. rodentium* infection (1-5×10^8^ *C. rodentium* CFU per mouse). WT naïve, n = 3; (S)WT naïve, n = 4; WT 1 day, n = 4; (S)WT 1 day, n = 4; WT 3 days, n = 5; (S)WT 3 days, n = Data is from one experiment. \**P* < 0.05, Student’s t-test.

Other cytokines, such as G-CSF and IP-10 showed a trend towards higher expression in (S)WT animals. The amount for other cytokines analyzed were similar between the two types of mice. In the colon, cytokine measurements showed greater variability between the two mouse groups (**Figure 6**). Although these differences did not reach statistical significance, G-CSF, IL-6, IL-7, IL-10, IL-17, IP-10, and TNFα tended to be higher in (S)WT mice baseline or post infection time points. (S)WT mice, however, presented lower IL-18 levels in comparison to WT counterparts.

**Figure 6.**
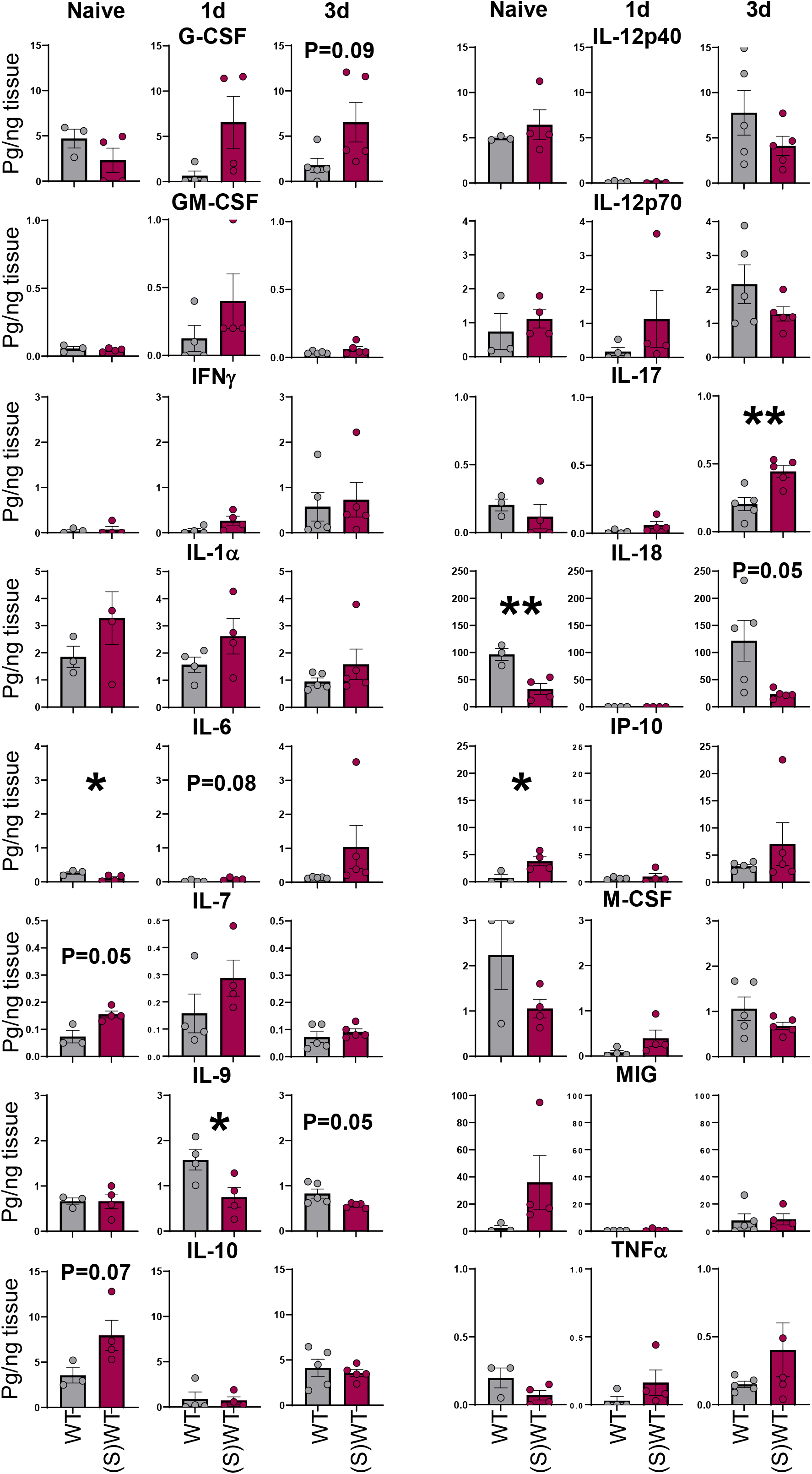
Detailed cytokine profile in the colon of C57BL/6 mice susceptible to *C. rodentium* infection. Data derived from the experiment in **Figure 4.** Individual cytokine expression in the cecum of WT and (S)WT mice at the indicated time points after *C. rodentium* infection (1-5×10^8^ *C. rodentium* CFU per mouse). WT naïve, n = 3; (S)WT naïve, n = 4; WT 1 day, n = 4; (S)WT 1 day, n = 4; WT 3 days, n = 5; (S)WT 3 days, n = 5. Data is from one experiment. \**P* < 0.05, Student’s t-test.

### Microbiota profile in susceptible C57BL/6 mice

We have previously reported that susceptible *Gzmb*^-/-^ mice to *C. rodentium* infection harbored an intestinal microbiota that is broadly similar to that of resistant counterparts, with the notable exception of reduced abundance of Turibacterales.^15^ Although these findings suggested that a related microbiota signature might also be present in the newly derived susceptible wild-type C57BL/6 [(S)WT] line, caution needs to be taken because loss of granzyme B could itself influence microbiota composition, including through indirect effects on bacterial survival or host-microbe interactions. We therefore analyzed the intestinal microbiota (stool samples) in 6-week-old WT and (S)WT mice. Alpha rarefaction analysis indicated that sequencing depth was sufficient to capture the microbial diversity present in both groups, and that the richness observed in (S)WT mice was slightly higher than in WT mice (**Figure 7A**). Unweighted UniFrac analysis showed strong separation between WT and (S)WT samples, indicating differences in community membership across the two groups of mice (**Figure 7B**). Weighted UniFrac analysis similarly demonstrated separation between the microbial communities present in WT and (S)WT (**Figure 7C**), suggesting that in addition to taxa differences, their relative abundances differed.

**Figure 7.**
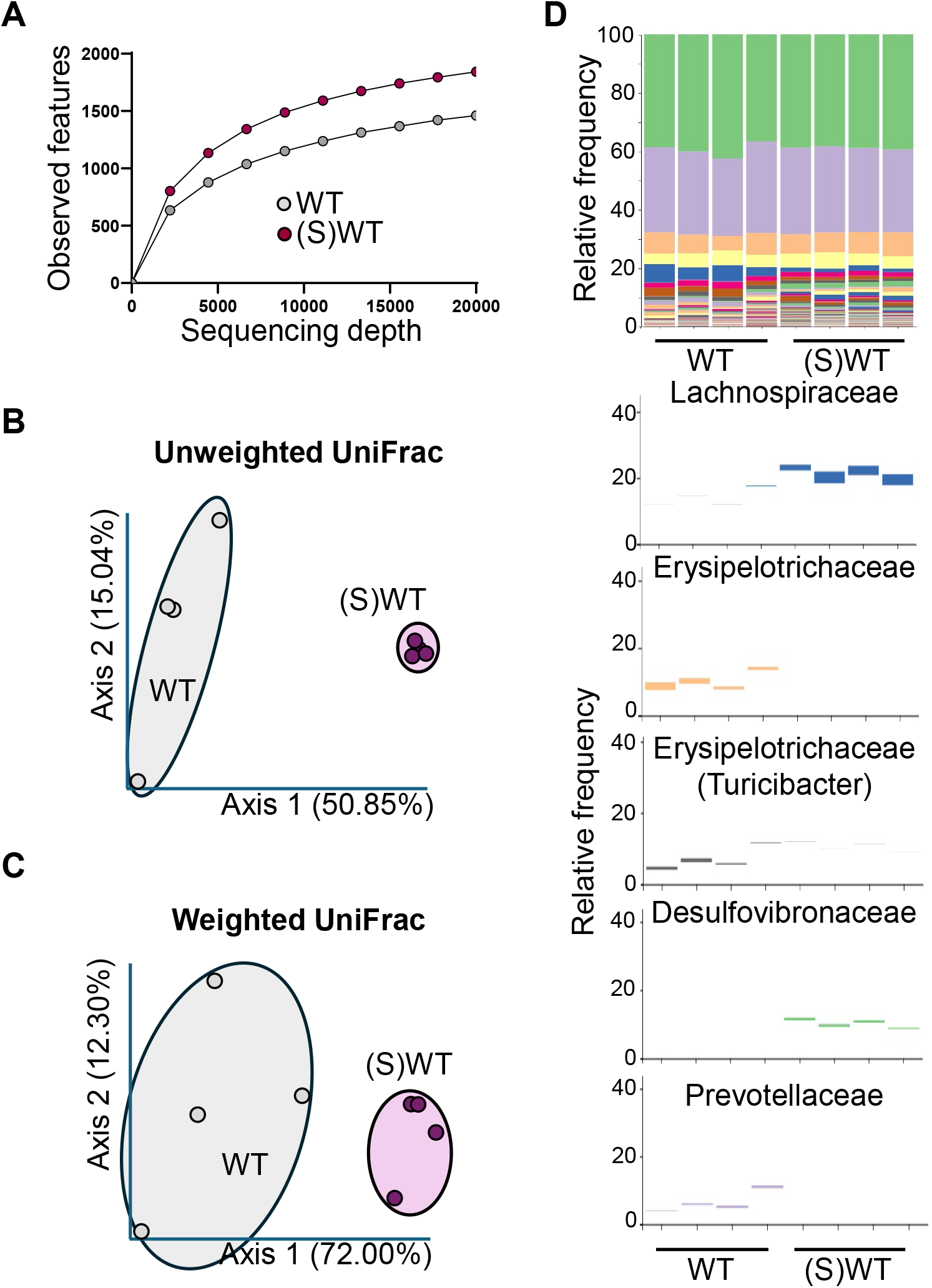
Stool microbiota analysis from C57BL/6 mice susceptible to *C. rodentium* infection. 16S rRNA DNA sequencing derived from stool samples of WT and (S)WT mice. (A) Alpha rarefaction analysis; (B) and (C) unweighted and weighted Emperor UniFrac analysis. (D) Taxonomic analysis. Top shows overall taxa between two groups; bottom, selected individual groups. WT naïve, n = 4; (S)WT naïve, n = 4. Data is representative of one experiment.

Taxonomic analysis showed similar abundance frequencies across major bacterial families (**Figure 7D**); however, families, such as Lachnospiraceae and Desulfovibronaceae were more prominent in the intestinal microbiota of (S)WT mice. The inverse was observed with other families, such as Erysipelotrichaceae, which were more abundant in the intestinal microbiota of WT mice. Of note, *Turicibacter* is grouped into Erysipelotrichaceae, indicating lower abundance of these bacteria in (S)WT mice. Together, these findings indicate that susceptible C57BL/6 mice harbor a distinct microbiota composition that parallels key features previously observed in susceptible *Gzmb*^-/-^ mice.

## Discussion

Animal models remain indispensable for defining infectious disease pathogenesis. Although animal models may not fully recapitulate what happens in humans, their use have yielded invaluable insight into mechanisms that can be targeted as potential interventions. In the case of human enteropathogenic and enterohemorrhagic *Escherichia coli* (EPEC and EHEC, respectively), efficient colonization of mice generally requires prior antibiotics treatment,^21^ a procedure that disrupts the intestinal microbiota and alters significant alterations to the intestinal microenvironment. *Citrobacter rodentium*, a natural intestinal pathogen of mice that produces intestinal disease has become one of the most important model for human EPEC and EHEC pathogens.^4^ Unlike EPEC and EHEC, *C. rodentium* infection does not require prior treatment with antibiotics, and a single inoculum administered by oral gavage or contaminated chow is sufficient to establish infection. The relevance of this model lies in its shared pathogenic features with human EPEC and EHEC pathogens, such as its intimate adherence to epithelial cells, effacement of microvilli, and actin-rich pedestal formation.^22^ Importantly, the lesions are mediated by conserved mechanisms, including a type III secretion system (T3SS) encoded by the locus of enterocyte effacement (LEE), similar to the human pathogens. However, one limitation of the model is that C57BL/6 mice, one of the most widely used laboratory strains, typically develop mild or no disease/pathology after infection with *C. rodentium*.^23,24^ Although studies using genetically altered C57BL/6 mice have provided important insights into host pathways that regulate susceptibility,^11,25-28^ they do not address severe disease in a non-genetically modified host.

Here, we report the derivation of a novel C57BL/6 line of mice with increased susceptibility to *C. rodentium* infection. This phenotype was first observed in granzyme B-deficient mice and it was thought that increased disease incidence and severity was caused by the absence of this enzyme.^11^ Although part of the phenotype was associated with granzyme B deficiency, we later concluded that the microbiota, in particular Turicibacterales, were responsible for part of the observed phenotype.^15^ As reported herein, when the susceptibility-induced microbiota was introduced into C57BL/6 WT mice, these mice developed severe infection characterized by wasting, loose stools, diarrhea, rectal bleeding, and severe cecal and colonic inflammation, which are rarely observed in conventional C57BL/6 WT mice.

A striking feature of the phenotype was its early onset. Susceptible mice began losing weight 24 hours post-infection indicating a cecal involvement during the first days after infection. Indeed, at day 3, cecum colonization and pathology was greater in (S)WT mice than in conventional WT animals (**Figure 1C**). These results indicate a potential defect in innate immune responses in these mice. Many innate immune cells are present in the intraepithelial lymphocyte (IEL) compartment: a group of diverse lymphoid cells intimately associated with intestinal epithelial cells.^29^ Their anatomical location allows IEL to rapidly respond against pathogens and other invaders. One-day post-infection, conventional WT mice showed increased IEL cellularity (**Figure 2B**), which serves as a surrogate parameter for IEL activation and responsiveness. This initial immune response may be of relevance to quench the infection and prevent development of a chronic disease. (S)WT mice, on the other hand, have reduced IEL cellularity (activation) suggesting a failure to induce an adequate innate immune response. It is noteworthy that the “activated” IEL observed in conventional WT mice corresponds to a variety of lymphocytes. For example, TCRγδ cells are known to move around intestinal epithelial cells to detect potential danger signals^30^ and induce tissue recovery;^31^ other cells, like iCD8α cells, capture and process antigen, and promote survival of other IEL populations;^32-34^ and TCRαβ^+^CD8αβ^+^ IEL represent *bona fide* tissue resident memory cells.^35^ Thus, the initial immune response in the cecum directed against *C. rodentium* is complex requiring the participation of different IEL subpopulations.

We hypothesized that an increase in cecal IEL cellularity in infected WT mice would correlate with higher cytokine levels in comparison to (S)WT mice. However, most of the cytokines analyzed, with the exception of G-CSF and IL-18, showed no expression differences between groups. The possibility exists that other cytokines, not covered in our panel, are differentially expressed among infected mouse groups. Or the increased levels of G-CSF and IL-18 in infected (S)WT mice may be sufficient to quench cellular proliferation. Lastly, it is also possible that there is not a direct correlation between cytokine expression and cecal IEL cellularity.

Microbiota profiling demonstrated clear separation between WT and (S)WT mice, supporting the conclusion that susceptibility is associated with a distinct intestinal microbial community. This underscores that the precautions we have taken to prevent cross-microbiota contamination between the two groups were sufficient to maintain both the microbiota configuration and the disease phenotype over time (**Figure 7**). In our previous report, we showed that the intestinal microbial communities between susceptible granzyme B-deficient and WT mice differed in the frequencies of Turicibacterales.^15^ Similarly, (S)WT mice showed decreased frequencies of Erysipelotrichceae, the taxonomic family for *Turicibacter. Turicibacter* appears to confer partial protection against severe *C. rodentium* infection,^15^ suggesting that the same phenotype observed in (S)WT may be due to decreased levels of this commensal. This is a current avenue of research our laboratory is undertaking.

How was this phenotype acquired? We originally obtained granzyme B-deficient mice from our collaborator, Dr. Xuefan Cao. We believe the susceptibility phenotype spontaneously arose and due to our stringent mouse husbandry protocols, the phenotype stabilized. We also kept susceptible granzyme B-deficient mice breeding as an independent line, i.e., without crossing them with conventional WT mice to generate granzyme B^-/-^ and control granzyme B^+/-^ littermates. Had the mutant line been routinely crossed to WT mice to generate littermate controls from the outset, the susceptibility phenotype may have been lost through microbiota normalization, as resistance is transmissible by cohousing.^15^ This observation should serve as a cautionary tale: although littermate-controlled breeding strategies are essential for rigorous genetic studies, they may also mask phenotypes that depend on transmissible microbial communities.

This susceptible C57BL/6 model represents a novel model for the study of EPEC and EHEC pathogens. The advantages of this model follow: a) discreet intestinal microbiota differences within the context of complex communities, avoiding the use of techniques to modify the microbiota; b) stability and reproducibility; c) the ‘susceptible’ microbiota can be stored, and transplanted into germ-free mice; and d) provides an opportunity to study how severe disease occurs without genetic manipulation.

Therefore, we believe this model will be useful for mechanistical studies related to the pathology of human EPEC and EHEC microorganisms.

## Materials and Methods

### Mice

C57BL/6 mice were originally purchased from The Jackson Laboratory (000664) and have been maintained in our colony for several years. Germ-free male was purchased from The Charles River Laboratories. Upon arrival, mouse was removed from isolating transport container and immediately set in a cage with two susceptible *Gzmb*^-/-^ female mice, originally provided by Dr. Xuefan Cao. The offspring of this breeding setup was bred among themselves to generate *Gzmb*^+/+^ mice, referred to as susceptible wild type mice, (S)WT. To prevent microbiome transfer between WT and (S)WT mice, cage husbandry was performed by members of our laboratory. Biosafety cabinet and all needed utensils were disinfected with MB-10 prior to handling mice.

Because resistance phenotype is transferable, (S)WT mice were always handled before WT mice. All mice were between 6 and 7 weeks of age at the time of *C. rodentium* infection. Male and female mice were used for all experiments.

### Citrobacter rodentium infection

Six- to 7-week-old mice were infected with *C. rodentium* (strain DBS100, ATCC 51459) as previously described.^15^ Briefly, bacteria colonies grown in McConkey plates, were picked and placed in 40ml of LB media (3 colonies per 40ml), and incubated overnight without shaking at 37°C. The next day, 8ml of culture were transferred into 32ml of fresh LB media and cultured for 4 hours with shaking at 37°C. OD_600_ was determined after the incubation period. We have determined that 1 OD equals approximately 1×10^9^ CFU. Bacteria was pelleted and resuspended in fresh LB media at 1-5×10^8^ CFU per 100μl. Mice were orally gavaged with 100μl containing 1-5×10^8^ CFU. Leftover inoculum was plated onto McConkey plates to confirm doses.

Starting weight was determined prior to gavage. Mice were weighed daily for the course of the experiment. At the same time, mice were monitored for the following signs of disease: irritated rectum (1 point), loose stool (1 point), rectal bleeding (1 point), diarrhea (1 point), hunched posture (1 point), scruffiness (2 points), and moribund (3 points). Mice with less than 0.5g of starting weight were considered as wasting, and 1 point was scored and included in disease score.

### Citrobacter rodentium colonization

At the indicated time points, ∼0.5cm section of the lower part of the cecum and the distal colon were dissected, weighed, placed in 0.5ml of LB and homogenized. After serial dilution, samples were placed on McConkey plates to determine CFU per gram of tissue.

### Citrobacter rodentium associated pathology

Swiss rolls from the indicated samples were fixed in 10% formalin buffered solution and then processed by the Translational Pathology Shared Resource Core. Pathology was performed blindly by Dr. Blanca Piazuelo. Score is based on inflammation, epithelial injury, hyperplasia, and architectural damage.

### Flow cytometric analysis of cecal lymphocytes

IEL and lamina propria cells were isolated as previously described.^32^ Briefly, after flushing the intestinal contents with cold HBSS and removing excess mucus, cecum was cut into small pieces (∼1 cm long) and shaken for 45 minutes at 37°C in HBSS supplemented with 5% fetal bovine serum and 2mM EDTA. Supernatants were recovered and IEL isolated using a discontinuous 40/70% Percoll (GE Healthcare Life Science) gradient. To obtain lamina propria lymphocytes, cecum tissue was recovered after IEL isolation and digested with collagenase (187.5 U/ml; Sigma-Aldrich) and DNase I (0.6 U/ml; Sigma-Aldrich). Cells were isolated using a discontinuous 40/70% Percoll gradient.

### Flow cytometry

The following fluorochrome-coupled anti-mouse antibodies were used: CD4 FITC, clone GK 1.5 (Tonbo). CD8α APC-H7, clone 53-6.7 (Tonbo). CD8β PE, clone YTS156.7.7 (Biolegend). CD45 PE-Cy7, clone 30-F11 (BD Biosciences). Ghost 510, viability (Tonbo). TCRβ PerCP-Cy5.5, clone H57-597 (Tonbo). TCRγ APC, clone GL3 (Invitrogen). All stained samples were acquired using a BD FACSCantoII, four laser Fortessa, or Five-laser LSR II flow cytometers (BD Biosciences), and data were analyzed using FlowJo software (Tree Star). Cell staining was performed following conventional techniques. Sample acquisition was performed at the Flow Cytometry Shared Resource at Vanderbilt University Medical Center.

### Cytokine analysis

Cecum and colon were dissected at the indicated time points, weighed and placed in 0.5ml of sterile PBS. Samples were homogenized with a mechanical tissue homogenizer and disposable 7mm generators (Fisher), followed by centrifugation to clear the supernatant from debris. Samples were stored at -70°C until further use. Cytokine Luminex profile was performed by Eve Technologies (Calgary, AB, Canada). Data was analyzed using Graph Prism.

### Intestinal microbiota analysis

Stool samples were collected from 6-week-old WT and (S)WT mice and quickly placed on ice for transport. Stool was stored at -70°C until further use. DNA was extracted using a DNeasy PowerSoil Pro Kit (QIAGEN), following the manufacturer’s instructions. Extracted DNA concentration was measured using Nanodrop 2000 (ThermoFisher Scientific). The V3 and V4 hypervariable regions of the 16SrRNA gene were sequenced using 2 × 300 paired-end-sequencing on the Illumina Miseq sequencing platform (Illumina) at VANTAGE (Vanderbilt University Medical Center). Primers 341F (CCTACGGGNGGCWGCAG) and 805R (GACTACHVGGGTATCTAATCC) were used for the V3-V4 region. The QIIME2 pipeline (version 3.6.11) was used to process and filter demultiplexed sequence reads.

Sequences were clustered using Deblur (2020.8.0) prior to alignment using QIIME2. OTU taxonomy was determined using a naive Bayesean classifier trained toward the GreenGenes 99% reference database (13_8). The OTU table was rarified to an even depth of references per sample prior to generation of taxonomy barplots using QIIME2view.

## Acknowledgments

This work was supported by R211AI187749 (W.H.). We acknowledge the Translational Pathology Shared Resource supported by NCI/NIH Cancer Center Support Grand P30CA068485. Flow Cytometry experiments were performed in the VMC Flow Cytometry Shared Resource, supported by the Vanderbilt Ingram Cancer Center (P30CA68485) and the Vanderbilt Digestive Disease Research Center (DK058404). Histopathology was performed by the Tissue Morphology subcore of the VDDRC (DK058404). 16S RNA DNA sequencing was performed at the Vanderbilt Technologies for Advance Genomics.

